# Alterations in sleep state boundaries and sleep dynamics following acute total and chronic partial sleep loss: a state space model exploration

**DOI:** 10.64898/2026.06.21.732147

**Authors:** Stefan Reutimann, Lukas Imbach, Zoé Burkhard, Christian R. Baumann, Angelina Maric

## Abstract

Chronic partial and acute total sleep loss have a distinct impact on sleep architecture. Namely, acute sleep deprivation primarily leads to a strong rebound of slow wave sleep, while chronic sleep restriction results in an increased propensity of REM sleep. The aim of this work was to examine whether these different effects would translate into quantifiable changes in sleep state boundaries and dynamics using a model-based method.

Besides conventional sleep stage scoring, we applied an EEG model (state space approach) for dynamic analysis of nocturnal EEG recordings in 14 healthy subjects under experimental chronic sleep restriction (last of 7 nights with 5 hours of time in bed) and after acute sleep deprivation (sleep following 40 hours of wakefulness), in comparison to baseline sleep.

Subjects under chronic sleep restriction revealed increased similarities in the frequency composition of REM sleep and wakefulness and thus, a decreased differentiation of state boundaries between the two behavioral states. Contrarily, acute sleep deprivation affected the spectral composition of NREM sleep. Only acute sleep deprivation resulted in more stable slow wave sleep.

Our explorative study confirmed that the distinct effects of increased REM sleep and slow wave sleep propensity following acute total and chronic partial sleep loss are reflected in differential changes of behavioral state boundaries and sleep dynamics. This suggests that these sleep structure characteristics are state dependent, which may allow using such measures in the future to track treatment effects in clinical populations characterized by sleep behavioral state dysregulation.

## Introduction

Experimental manipulations of sleep duration are widely used to model effects of sleep loss on cognition, behavior, metabolism, and sleep regulation.[1] The majority of past research applied acute sleep deprivation (ASD) protocols,[2] in which sleep is completely absent over a consecutive period.[1, 3] Such a total ASD is contrasted by more chronic, but partial forms of sleep loss, which resemble every day sleep loss more closely and may also relate better to sleep loss in a clinical context. For example, in a chronic sleep restriction (CSR) protocol, sleep opportunities are reduced over multiple, consecutive nights.[1, 3] These two paradigms have been shown to result in distinct changes of consequent sleep propensity: Following the complete sleep loss in ASD, subsequent sleep primarily displays a strong slow wave sleep (SWS) rebound[4, 5], as evident by increased portions of SWS,[6–10] enhanced slow wave activity and density[9, 11], together with decreased SWS latency[6, 8, 10]. The rebound in rapid-eye-movement sleep (REMS) is more limited and mostly evident after extensive ASD only[6–8, 10]. Contrarily, CSR results in increased REMS propensity, as evident by increased portions and decreased latency of REMS[2, 10, 12–15], while increases in SWS propensity have been found to be less pronounced[10, 13, 15]. These distinctive effects are attributed to a higher priority for compensation of SWS if all sleep had been absent on the one hand, as it is the case in ASD. On the other hand, curtailing sleep periods, as present in CSR, leads to a predominant loss of REMS and greater preservation of SWS, since earlier parts of a sleep period are typically rich in SWS, while REMS typically dominates later parts.[16]

While the sleep stage categorizations obtained by conventional sleep stage scoring, provide important information on the global architecture of sleep, further quantitative information may be gained by spectral analysis of the EEG signal. The spectral power density in the delta range for example, is typically used to quantify the intensity of SWS and therefore sleep depth.[17] However, this approach reveals only limited information on sleep dynamics and sleep state boundaries. A recently introduced method, which allows reliable quantitative assessments of such aspects, is the state space analysis (SSA) approach.[18] Briefly, in SSA, sleep is represented by a two-dimensional space, derived from its spectral characteristics. Spectral compositions can be compared across sleep stages based on the resulting cluster arrangements and state stability can be derived from the distance in space between consecutive epochs.[18]

By applying the SSA in clinical populations, it was possible to quantify pathological alterations in sleep state boundaries and sleep dynamics.[18, 19] Importantly, such measures may even qualify as a diagnostic tool, for example in Parkinson’s disease or Narcolepsy.[18, 19] However, up to date it is unknown, whether sleep characteristics as derived from SSA, reflect stable (pathological) predispositions or if these features can be modulated by sleep regulation mechanisms.

Here, we explored SSA for assessing changes in sleep macrostructure and dynamics induced by CSR and ASD in healthy subjects. While previous work described differential effects of CSR and ASD on sleep architecture and spectral EEG characteristics, it remains unclear whether these distinct effects are also reflected in quantitative measures of behavioral state boundaries and sleep dynamics as derived from SSA. Based on the increased REMS propensity after CSR, and the increased SWS propensity after ASD, we expected to find differences in parameters relating to REMS macrostructure and dynamics in CSR and SWS in ASD, respectively.

## Methods

### Study population and protocols

We included nocturnal EEG data of 14 young, healthy, male individuals (21.9 +-3.0) years, mean (+/- SD) who participated in a previously published experimental sleep restriction and sleep deprivation study.[10] Subjects had been recruited from a student population and had been required to have habitual sleep durations between 7 and 9 hours. The counter-balanced, cross-over study design included an ASD of 40 hours and a CSR with a limited sleep opportunity of five hours per night for seven consecutive nights. Both manipulations were preceded by one week of regular sleep rhythm and the wash-out period between conditions was at least two weeks. Napping during daytime was forbidden and participants were asked to refrain from medication, alcohol, and caffeine intake. In all subjects’ sleep was recorded in the laboratory at baseline (BSL, i.e., after one week of regular sleep), after ASD, and during the last 3 nights of CSR using a high-density EEG net (Sensor Net for long-term monitoring; Electrical Geodesics Inc., Eugene, OR) with a sampling rate of 500 Hz. A more detailed description on the study design and procedures is provided elsewhere.[10] The study was approved by the local ethics committee, and all participants gave written informed consent. Three different nights were included in the current analysis: the BSL night, the last (seventh) night of CSR, and the recovery night after ASD.

### EEG preprocessing

As previously described,[10] the EEG was filtered (0.5 Hz high-pass, 40 Hz low-pass filter), bad-quality channels and artefacts visually and semi-automatically removed, and re-referenced to the average of all electrodes (average reference). For the current analysis, all data of every night was shortened to 295 minutes, as this corresponded to the minimal duration available in all recordings.

### Sleep Architecture

Sleep scoring was performed according to standard criteria adapted to be applied to 20-second epochs.[10] Sleep onset was defined as the first occurrence of stage N1 (or in theory, any other sleep stage reached first). N2, N3 and stage R latencies were calculated from sleep onset. Sleep efficiency was defined as the ratio of total sleep time relative to the time analyzed. Fragmentation of individual sleep stages was calculated as the percentage of all epochs of a given sleep stage that were immediately followed by an epoch of any other stage, relative to the total number of epochs in the sleep stage.

### State space analysis

All EEG analyses were executed using MATLAB (The Mathworks Inc., Natick, MA, Version R2018b).

For construction of the EEG state space, the EEG data was divided into epochs of five seconds length, labelled with the corresponding sleep stage. The mean spectral power density of the EEG signal over six standard electrode positions (F3, F4, C3, C4, O1, O2; average reference) was calculated by Welch’s power spectral density estimate using a 5-second-Hamming window. Subsequently, two spectral ratios (Ratio 1 = (8.6 to 19.3 Hz)/(1.0 to 10.9 Hz) and Ratio 2 = (11.5 to 20.3 Hz)/(17.9 to 31.5 Hz)) were computed for each epoch, based on the composition of the power spectral density, as described previously[20]. The resulting time series was smoothed by a 50-Hann window. The logarithm of Ratio 1 determined the x-position and the logarithm of Ratio 2 the position in the y-direction of the corresponding epoch in space.

The velocity in state space was defined as the Euclidean distance in space between two subsequent epochs, divided by the elapsed time between them. Thus, a low velocity demonstrated little spatial change between two consecutive epochs, indicating little alteration in spectral composition and consequently, a stable, consolidated sleep period. A velocity cutoff was used to determine stable and unstable sleep, as defined previously[18]: A velocity of < 0.01 indicated sleep stage clusters, i.e., stable states, while a velocity of > 0.01 indicated trajectories, i.e., unstable states. The number of stable epochs per sleep stage was assessed as a measure of sleep stage stability.

For cluster analysis, the position of the centroid of each sleep stage cluster was calculated by taking the mean of the positions of all stable epochs in the corresponding sleep stage. The distance between each pair of centroids was calculated to determine the differences in the spectral compositions of the sleep stages.

To verify the concept of cluster centroids, and to analyze spatial distribution of the different sleep stages, the point density of the stable epochs of each sleep stage was further evaluated in x- and in y-direction using a one-dimensional probability approach, based on the normal kernel function. Here the clusters of stage N2 and N3 combined to one N2/N3 cluster, due to the closeness of the spectral characteristics of the two stages resulting in overlaps of the two probability density functions. The peak-to-peak distance was assessed to detect similarities and differences in the point densities of the different sleep stage clusters.

Additionally, the one-dimensional point densities in x- and y-direction were calculated including unstable and stable epochs combined, to further include trajectories between and within the sleep stages into analysis. The overlap areas of the point densities of each two sleep stages were determined by calculating the cumulative integral of the intersection of the two corresponding point densities.

To derive the number of transitional periods between the different sleep stages, a two-dimensional probability density approach was executed. First, the multivariate normal distribution of each sleep stage was calculated. Transitions were determined by assigning each epoch to the sleep phase, which revealed the highest point density value at the position of the epoch. Each pair of subsequent epochs leaving a sleep stage and ending in another was counted as a transitional period. This was done for every possible pair sleep stage in both ways (R-N2/3, W-R, W-N2/N3).

### Statistics

Statistical analysis was executed using SPSS software program (IBM Corp., Armonk, NY, Version 26.0, 2019). Since normality could not be assumed for all variables (p<0.05 in Shapiro-Wilk test of normality) Wilcoxon signed-rank tests were applied throughout for pairwise comparisons between conditions. These analyses included conventional sleep parameters, positions and distances of cluster centroids, peak-to-peak distances, overlap areas, the number and ratio of stable epochs, as well as measures of behavioral state fragmentation and transitional periods between behavioral states. Bonferroni correction was applied to all p-values to correct for multiple comparisons across the three conditions (i.e., multiplication of p-values by 3).

## Results

### Sleep architecture

The conventional sleep scoring revealed changes in sleep architecture after CSR and ASD that were consistent with increased sleep pressure due to the induced sleep loss (Table 1). Namely, we observed a lower proportion of lighter sleep stages (N1, N2), shorter sleep latency, and an increased sleep efficiency – with some of these effects tending to be stronger after ASD than during CSR.

**Table 1:**
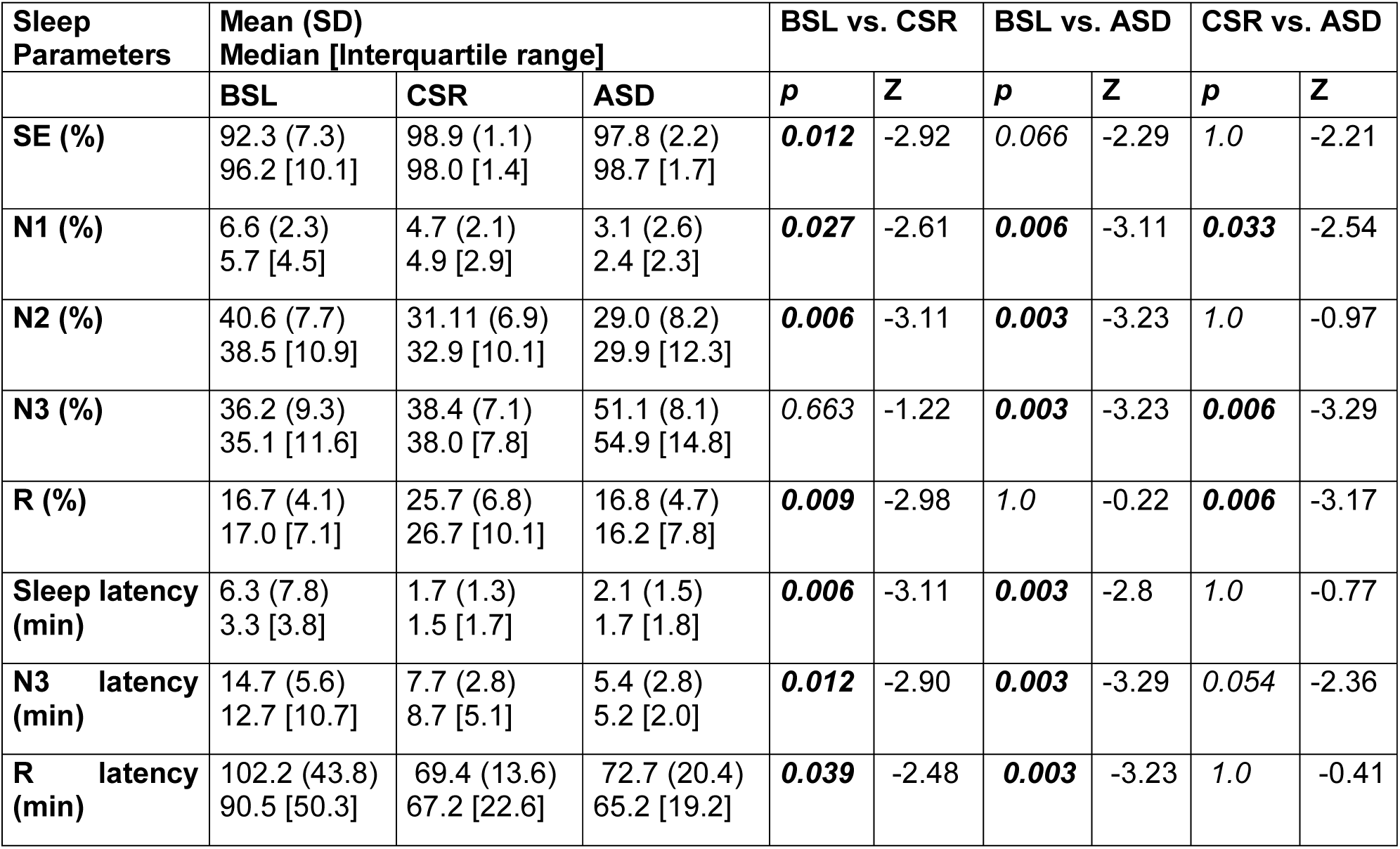
Sleep parameters as derived from conventional sleep scoring. Data is shown as mean (+/- SD) and median [interquartile range] with corresponding p- and z-values resulting from the Wilcoxon signed-rank test. Bonferroni correction was applied to all p-values. Bold values mark significant differences between conditions. BSL: Baseline sleep; CSR: Chronic sleep restriction; ASD: Acute sleep deprivation; SE: Sleep efficiency; N1, N2, N3, and R refer to sleep stages as defined in the AASM Scoring Manual.[36]

Latencies to stages N3 and R were reduced in both conditions. However, the percentage of N3 was significantly higher only after ASD, but not during CSR. Contrarily, the proportion of stage R sleep was significantly higher in CSR, but not after ASD. These results confirm that also in our sample CSR predominantly increased REMS propensity, whereas ASD mainly increased SWS propensity.

### Spectral composition and distinction of behavioral states

To assess the overlap of spectral features between the different behavioral states, relations of different clusters in space were evaluated. Figure 1 shows state space clusters of stable epochs in all three conditions of one representative subject. When comparing the cluster centroid positions of the different behavioral states we found significant displacements of several clusters after experimental sleep loss (Table 2). ASD resulted in a significant disarrangement of the N2 and N3 clusters, mostly in the y-dimension, primarily based on a change in the frequency band of 11.5 to 20.3 Hz. During CSR only the N2, but not the N3 cluster, tended to be shifted as well. Contrarily, we found a significant displacement of the stage R cluster in the x-dimension during CSR, but not after ASD. This displacement was based on reductions in both frequency bands, namely the band between 8.6 to 19.3 Hz and 1.0 to 10.9 Hz, with the latter band showing a stronger reduction. The shift of the stage R cluster during CSR resulted in an increased similarity in the spectral compositions of stage W and stage R, as evident by a significantly smaller distance of the cluster centroids (Table 3). This is also evident in the CSR state space plot example in Figure 1B compared to the BSL state space plot in Figure 1A. A significantly lowered peak-to-peak distance of the point density estimates (again including only stable epochs; Figure 2, Table 4) and a significantly larger overlap area of the point density functions (including also unstable epochs; Table 5), further confirmed this increase in similarity of the spectral characteristics between stage W and stage R during CSR. Nevertheless, these findings need to be considered in the context of the comparatively low number of wake epochs after the sleep deprivation conditions (see Table 6).

**Figure 1:**
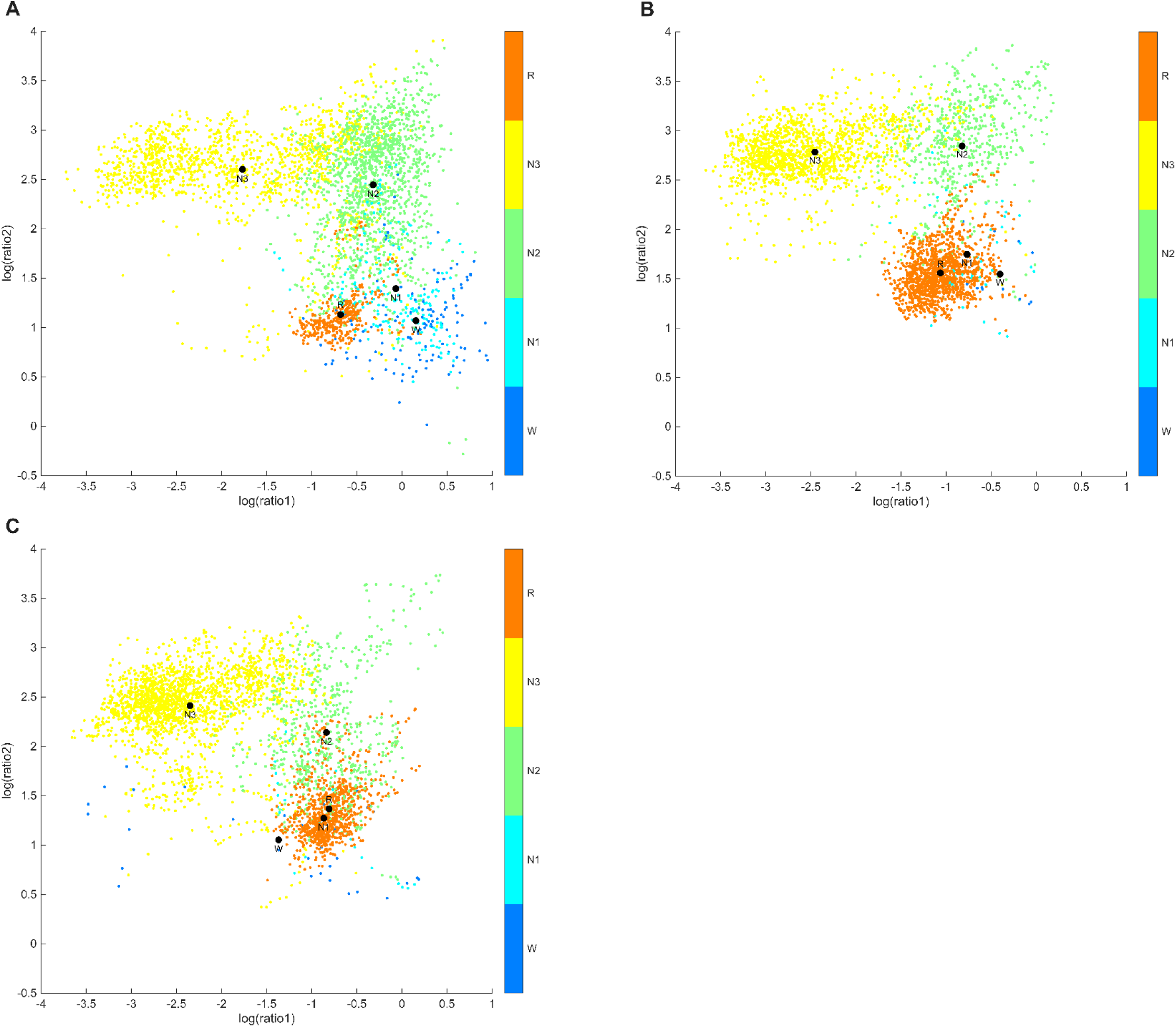
Representative state space plots of one subject **(A)** during baseline (BSL), **(B)** during chronic sleep restriction (CSR), and **(C)** after acute sleep deprivation (ASD), smoothed with 10-Hann window. Cluster centroids are presented as black circles. Vigilance stages (W, N1, N2, N3, R) are color coded.

**Figure 2:**
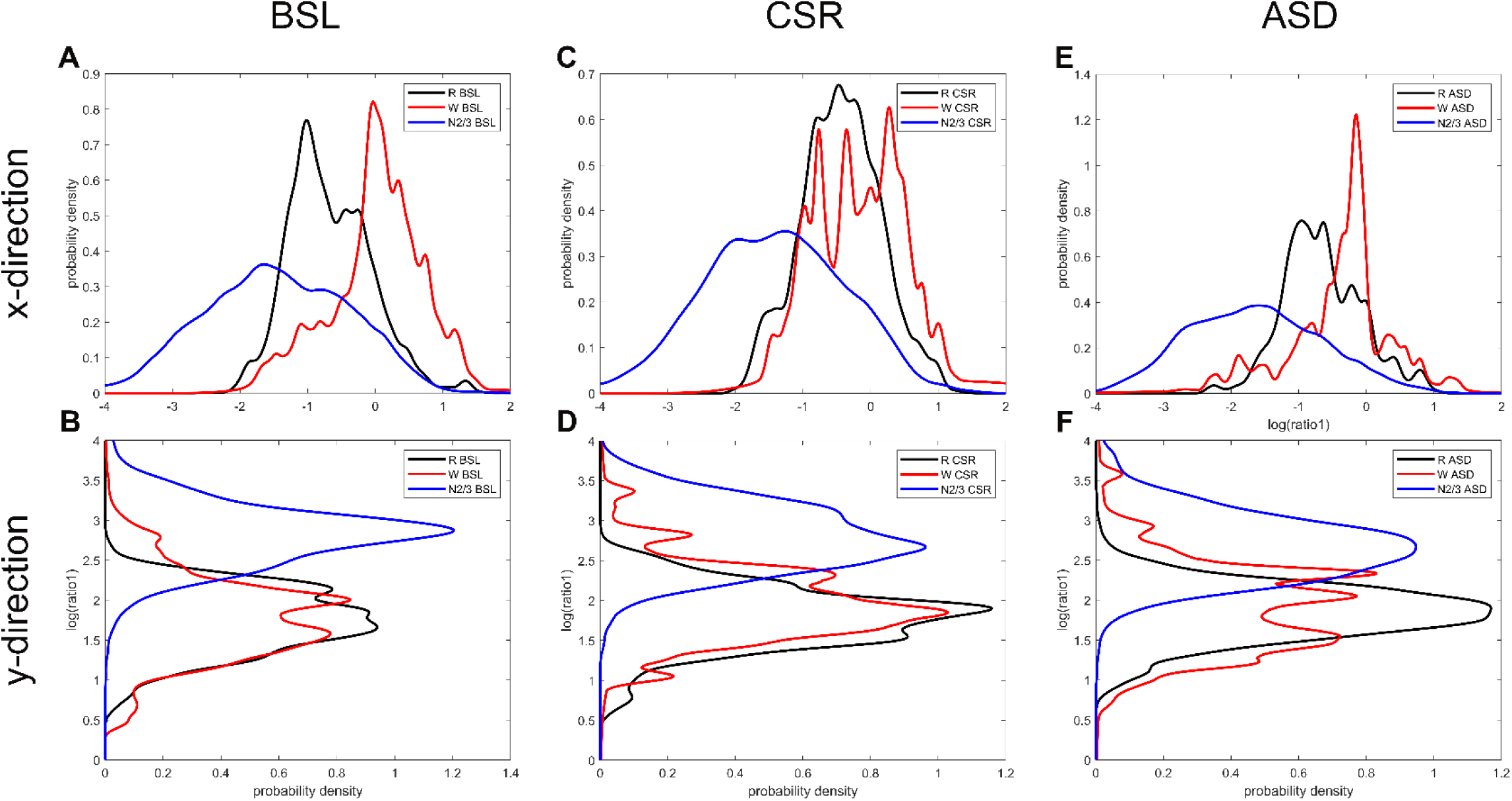
Point density curves of stage R, Wakefulness (W) and stage N2/3 of stable states **(A)** at baseline (BSL) in x-direction, **(B)** and in y-direction, **(C)** after chronic sleep restriction (CSR) in x-direction, **(D)** and in y-direction, **(E)** after acute sleep deprivation (ASD) in x-direction and **(F)** in y-direction. Vigilance stages (W, N2/3, R) are color coded and refer to sleep stages as defined in the AASM Scoring Manual (N2/3 are stages N2 and N3 combined) [36].

**Table 2:** Analysis of position of centroids of the different behavioral states. Data is shown as mean (+/- SD) and median [interquartile range] with corresponding p- and z-values resulting from the Wilcoxon signed-rank test. Bonferroni correction was applied to all p-values. Bold values mark significant differences between conditions. BSL: Baseline sleep; CSR: Chronic sleep restriction; ASD: Acute sleep deprivation; N1, N2, N3, and R refer to sleep stages as defined in the AASM Scoring Manual.[36]

|  | BSL | CSR | ASD | BSL vs. CSR |  | BSL vs. ASD |  | CSR vs. ASD |  |
| --- | --- | --- | --- | --- | --- | --- | --- | --- | --- |
|  |  |  |  | *p* | *Z* | *p* | *Z* | *p* | *Z* |
| **Position centroids** |  |  |  |  |  |  |  |  |  |
| W x | -0.01 (0.40)<br>0.01 [0.70] | -0.16 (0.29)<br>-0.19 [0.45] | -0.40 (0.64)<br>-0.30 [1.01] | 0.519 | -1.363 | 0.117 | -2.062 | 0.165 | -1.922 |
| W y | 1.87 (0.36)<br>1.89 [0.59] | 1.98 (0.32)<br>1.90 [0.40] | 1.96 (0.46)<br>1.88 [0.66] | 1.000 | -0.594 | 1.000 | -0.874 | 1.000 | -0.454 |
| N1 x | -0.44 (0.49)<br>-0.50 [0.75] | -0.30 (0.37)<br>-0.34 [0.57] | -0.48 (0.64)<br>-0.64 [0.60] | 0.594 | -1.287 | 1.000 | -0.659 | 0.327 | -1.601 |
| N1 y | 2.01 (0.41)<br>2.04 [0.48] | 2.10 (0.28)<br>2.11 [0.38] | 2.11 (0.45)<br>2.16 [0.68] | 0.663 | -1.224 | 0.900 | -1.036 | 1.000 | -0.596 |
| N2 x | -0.89 (0.49)<br>-0.96 [0.71] | -0.61 (0.44)<br>-0.64 [0.50] | -0.76 (0.46)<br>-0.82 [0.80] | 0.066 | -2.291 | 0.090 | -2.166 | 0.372 | -1.538 |
| N2 y | 2.72 (0.28)<br>2.67 [0.50] | 2.69 (0.31)<br>2.59 [0.61] | 2.50 (0.26)<br>2.44 [0.42] | 1.000 | -0.785 | **0.006** | -3.170 | **0.006** | -3.045 |
| N3 x | -2.05 (0.64)<br>-2.15 [0.50] | -1.91 (0.56)<br>-1.89 [0.79] | -1.99 (0.51)<br>-2.02 [0.61] | 0.474 | -1.412 | 1.000 | -0.722 | 0.598 | -1.287 |
| N3 y | 2.83 (0.22)<br>2.75 [0.39] | 2.84 (0.21)<br>2.83 [0.36] | 2.73 (0.22)<br>2.68 [0.40] | 1.000 | -0.345 | **0.006** | -3.107 | 0.078 | -2.229 |
| R x | -0.67 (0.43)<br>-0.78 [0.62] | -0.44 (0.34)<br>-0.44 [0.53] | -0.58 (0.45)<br>-0.61 [0.73] | **0.033** | -2.542 | 0.531 | -1.350 | 0.327 | -1.601 |
| R y | 1.74 (0.26)<br>1.76 [0.38] | 1.79 (0.28)<br>1.80 [0.32] | 1.81 (0.29)<br>1.82 [0.39] | 0.394 | -1.287 | 0.816 | -1.099 | 1.000 | -0.220 |

**Table 3:** Analysis of distance between the different centroids of the different behavioral states. Data is shown as mean (+/- SD) and median [interquartile range] with corresponding p- and z-values resulting from the Wilcoxon signed-rank test. Bonferroni correction was applied to all p-values. Bold values mark significant differences between conditions. BSL: Baseline sleep; CSR: Chronic sleep restriction; ASD: Acute sleep deprivation; N1, N2, N3, and R refer to sleep stages as defined in the AASM Scoring Manual.[36]

|  | BSL | CSR | ASD | BSL vs. CSR |  | BSL vs. ASD |  | CSR vs. ASD |  |
| --- | --- | --- | --- | --- | --- | --- | --- | --- | --- |
|  |  |  |  | *p* | *Z* | *p* | *Z* | *p* | *Z* |
| **Distance centroids** |  |  |  |  |  |  |  |  |  |
| W-R | 0.83 (0.27)<br>0.80 [0.43] | 0.53 (0.31)<br>0.46 [0.50] | 0.69 (0.37)<br>0.56 [0.38] | **0.021** | -2.691 | 0.348 | -1.572 | 1.000 | -0.943 |
| W-N2 | 1.31 (0.50)<br>1.33 [0.68] | 0.97 (0.40)<br>0.99 [0.52] | 0.91 (0.39)<br>0.80 [0.63] | **0.048** | -2.411 | 0.069 | -2.271 | 1.000 | -0.105 |
| W-N3 | 2.35 (0.56)<br>2.26 [1.11] | 2.05 (0.47)<br>2.12 [0.81] | 1.93 (0.56)<br>1.87 [0.63] | 0.084 | -2.201 | 0.225 | -1.782 | 1.000 | -0.314 |
| R-N2 | 1.08 (0.28)<br>1.05 [0.44] | 0.98 (0.25)<br>0.96 [0.35] | 0.77 (0.18)<br>0.77 [0.30] | 0.066 | -2.291 | **0.003** | -3.233 | **0.006** | -3.107 |
| R-N3 | 1.86 (0.43)<br>1.96 [0.56] | 1.88 (0.32)<br>1.87 [0.37] | 1.79 (0.32)<br>1.74 [0.41] | 1.000 | -0.408 | 1.000 | -0.847 | 0.123 | -2.040 |

**Table 4:** Analysis of peak-to-peak distance of the point density functions of the different behavioral states. Data is shown as mean (+/- SD) and median [interquartile range] with corresponding p- and z-values resulting from the Wilcoxon signed-rank test. Bonferroni correction was applied to all p-values. Bold values mark significant differences between conditions. BSL: Baseline sleep; CSR: Chronic sleep restriction; ASD: Acute sleep deprivation; N1, N2, N3, and R refer to sleep stages as defined in the AASM Scoring Manual.[36]

|  | BSL | CSR | ASD | BSL vs. CSR |  | BSL vs. ASD |  | CSR vs. ASD |  |
| --- | --- | --- | --- | --- | --- | --- | --- | --- | --- |
|  |  |  |  | *p* | *Z* | *p* | *Z* | *p* | *Z* |
| **Peak-to-peak distance** |  |  |  |  |  |  |  |  |  |
| W-R x | 0.72 (0.30)<br>0.64 [0.54] | 0.41 (0.28)<br>0.41 [0.46] | 0.62 (0.37)<br>0.52 [0.61] | **0.03** | -2.578 | 0.225 | -1.778 | 0.066 | -2.134 |
| W-R y | 0.35 (0.23)<br>0.29 [0.34] | 0.33 (0.30)<br>0.18 [0.51] | 0.39 (0.21)<br>0.29 [0.33] | 1.0 | -0.357 | 0.69 | -1.201 | 1.000 | -0.800 |
| W-N2/N3 x | 1.21 (0.58)<br>1.32 [1.04] | 1.69 (0.61)<br>1.54 [1.02] | 1.79 (0.74)<br>1.87 [1.16] | 0.297 | -1.647 | 0.078 | -2.223 | 1.000 | -0.800 |
| W-N2/N3 y | 0.95 (0.40)<br>1.03 [0.58] | 0.86 (0.36)<br>0.85 [0.54] | 0.76 (0.43)<br>0.53 [0.76] | 1.0 | -0.267 | 0.393 | -1.511 | 0.984 | -0.978 |
| R-N2/N3 x | 0.63 (0.43)<br>0.72 [0.79] | 1.55 (0.42)<br>1.38 [0.73] | 1.46 (0.60)<br>1.50 [1.04] | **0.003** | -3.296 | **0.003** | -3.296 | 0.993 | -0.973 |
| R-N2/N3 y | 1.14 (0.27)<br>1.13 [0.46] | 1.09 (0.25)<br>1.13 [0.29] | 0.95 (0.27)<br>0.94 [0.23] | 1.0 | -0.910 | **0.027** | -2.605 | **0.018** | -2.731 |

**Table 5:** Analysis of overlap areas of the point density functions of the different behavioral states. Data is shown as mean (+/- SD) and median [interquartile range] with corresponding p- and z-values resulting from the Wilcoxon signed-rank test. Bonferroni correction was applied to all p-values. Bold values mark significant differences between conditions. BSL: Baseline sleep; CSR: Chronic sleep restriction; ASD: Acute sleep deprivation; N1, N2, N3, and R refer to sleep stages as defined in the AASM Scoring Manual.[36]

|  | BSL | CSR | ASD | BSL vs. CSR |  | BSL vs. ASD |  | CSR vs. ASD |  |
| --- | --- | --- | --- | --- | --- | --- | --- | --- | --- |
|  |  |  |  | *p* | *Z* | *p* | *Z* | *p* | *Z* |
| **Overlap area** |  |  |  |  |  |  |  |  |  |
| W-R x | 0.27 (0.14)<br>0.27 [0.25] | 0.40 (0.15)<br>0.40 [0.25] | 0.30 (0.14)<br>0.31 [0.17] | **0.027** | -2.605 | 1.000 | -0.471 | 0.192 | -1.852 |
| W-R y | 0.48 (0.17)<br>0.49 [0.28] | 0.52 (0.14)<br>0.52 [0.15] | 0.43 (0.16)<br>0.45 [0.18] | 1.0 | -0.175 | 1.000 | -0.874 | 0.069 | -2.271 |
| W-N2/N3 x | 0.26 (0.13)<br>0.24 [0.25] | 0.33 (0.12)<br>0.32 [0.17] | 0.33 (0.20)<br>0.34 [0.24] | 0.252 | -1.726 | 0.474 | -1.412 | 1.000 | -0.534 |
| W-N2/N3 y | 0.18 (0.13)<br>0.15 [0.25] | 0.21 (0.12)<br>0.21 [0.13] | 0.21 (0.13)<br>0.20 [0.21] | 1.0 | -0.910 | 1.0 | -0.785 | 1.0 | -0.408 |
| R-N2/N3 x | 0.36 (0.13)<br>0.39 [0.23] | 0.31 (0.08)<br>0.31 [0.12] | 0.27 (0.09)<br>0.26 [0.14] | 0.531 | -1.350 | 0.066 | -2.291 | 0.090 | -2.166 |
| R-N2/N3 y | 0.09 (0.04)<br>0.08 [0.11] | 0.11 (0.07)<br>0.08 [0.53] | 0.16 (0.07)<br>0.15 [0.11] | 0.735 | -1.161 | **0.033** | -2.542 | **0.039** | -2.480 |

**Table 6:** Number of stable epochs and ratio of stable epochs relative to all epochs in each sleep stage. Data is shown as mean (+/- SD) and median [interquartile range] with corresponding p- and z-values resulting from the Wilcoxon signed-rank test. Bonferroni correction was applied to all p-values. Bold values mark significant differences between conditions. BSL: Baseline sleep; CSR: Chronic sleep restriction; ASD: Acute sleep deprivation; N1, N2, N3, and R refer to sleep stages as defined in the AASM Scoring Manual.[36]

|  | BSL | CSR | ASD | BSL vs. CSR |  | BSL vs. ASD |  | CSR vs. ASD |  |
| --- | --- | --- | --- | --- | --- | --- | --- | --- | --- |
|  |  |  |  | *p* | *Z* | *p* | *Z* | *p* | *Z* |
| Stable epochs (#) |  |  |  |  |  |  |  |  |  |
| W | 94.37 (109.81)<br>39.25 [138.71] | 10.95 (7.44)<br>9.25 [8.75] | 13.71 (16.84)<br>8.75 [14.96] | **0.006** | -3.107 | **0.033** | -2.542 | 1.0 | -0.345 |
| N1 | 55.45 (33.11)<br>51.17 [44.58] | 41.82 (35.06)<br>42.17 [55.42] | 22.18 (23.88)<br>15.58 [25.21] | 0.288 | -1.665 | **0.003** | -3.233 | **0.018** | -2.760 |
| N2 | 469.65 (154.25)<br>431.75 [238.58] | 385.88 (113.56)<br>368.08 [144.75] | 390.45 (133.67)<br>430.50 [227.71] | 0.192 | -1.852 | 0.501 | -1.381 | 1.0 | -0.157 |
| N3 | 460.51 (114.03)<br>416.08 [168.83] | 619.33 (127.42)<br>580.50 [165.92] | 871.39 (122.75)<br>873.17 [189.83] | **0.012** | -2.856 | **0.003** | -3.296 | **0.006** | -3.170 |
| R | 282.14 (89.87)<br>280.08 [192.17] | 469.95 (147.92)<br>429.50 [239.21] | 295.01 (111.86)<br>304.33 [159.33] | **0.012** | -2.856 | 1.0 | -0.094 | **0.006** | -3.107 |
| Ratio stable epochs |  |  |  |  |  |  |  |  |  |
| W | 0.298 (0.126)<br>0.254 [0.198] | 0.159 (0.071)<br>0.148 [0.099] | 0.158 (0.113)<br>0.132 [0.179] | **0.012** | -2.794 | **0.048** | -2.417 | 1.0 | -0.157 |
| N1 | 0.247 (0.096)<br>0.258 [0.141] | 0.223 (0.133)<br>0.239 [0.275] | 0.188 (0.107)<br>0.210 [0.152] | 1.0 | -0.471 | 0.105 | -2.103 | 0.531 | -1.350 |
| N2 | 0.349 (0.063)<br>0.351 [0.069] | 0.353 (0.051)<br>0.356 [0.064] | 0.382 (0.062)<br>0.392 [0.072] | 1.0 | -0.157 | 0.372 | -1.538 | 0.372 | -1.538 |
| N3 | 0.403 (0.092)<br>0.403 [0.124] | 0.469 (0.070)<br>0.454 [0.104] | 0.470 (0.054)<br>0.500 [0.073] | 0.057 | -2.354 | **0.006** | -3.170 | 0.372 | -1.538 |
| R | 0.517 (0.084)<br>0.521 [0.108] | 0.529 (0.093)<br>0.567 [0.147] | 0.499 (0.095)<br>0.517 [0.157] | 0.900 | -1.036 | 1.0 | -0.471 | 0.372 | -1.538 |

The above-mentioned stage N2 cluster centroid changes resulted on the one hand in a significantly smaller distance to stage R after ASD, with a corresponding trend during CSR, and a significantly smaller distance to stage W during CSR, with a corresponding trend after ASD on the other hand (Table 3). Interestingly, the point density estimate analysis revealed a significantly increased peak-to-peak distance between stage R and N2/N3 in x-direction after ASD and CSR, but a significantly lowered separation in y-direction after ASD only (Figure 2, Table 4). Along these lines, ASD led to a significant expansion of the overlap area between stage R and N2/N3 in y-direction, but a trend-like decrease of the overlap in x-direction (Table 5). Therefore, ASD affected state separation differently across the two dimensions of the state space: separation increased in x-direction but decreased in y-direction.

In summary, CSR predominantly affected features of the stage R cluster, which was not observed after ASD. This included a significant drift in the centroids’ x-axis position, lower distance between centroids of stages R and W, decreased peak-to-peak distance of stage R and stage W point density estimates in x-direction and increased overlap area of the two behavioral states which included not only stable but also unstable epochs. Together, these findings suggest a lower separation of the corresponding point densities and therefore a lower spectral distinction between R and W during CSR, particularly in x-direction. On the other hand, ASD significantly influenced the characteristics of N2 and N3 sleep, whereas CSR revealed a small but more restricted effect on these characteristics as well. Namely, ASD caused a significant displacement of the N2 and N3 clusters mainly in y-direction, significant changes in cluster distances, peak-to-peak distances and overlap areas of the N2/N3 sleep cluster relative to the stage R cluster. Thus, the point density analysis suggests that ASD did not simply reduce or increase state separation globally but rather altered the relation between behavioral states in a frequency-specific manner.

### Sleep dynamics

To assess the stability of behavioral states, we evaluated the number of stable epochs according to the velocity in the state space in each stage. We found a significant decrease of stable epochs in stage W, and an increase in stable stage N3 epochs after ASD and during CSR, with this increase in N3 being stronger after ASD compared to CSR (upper part Table 6). The absolute number of stable stage R epochs was only increased during CSR, but not after ASD. Interestingly, the changes in the number of stable epochs in stages W and N3 also translated into a change in the proportion of stable epochs relative to all epochs in these sleep stages, while this was not found for the number of stable epochs in stage R (lower part Table 6). This implies that the larger number of consolidated stage R epochs after CSR was not explained by an increased stability of REMS, but only by the increase in the total amount of REMS during CSR. In contrast, the stability of SWS after ASD was found to be increased (which tended to be the case during CSR as well).

These implications were further supported by assessing the corresponding sleep stage fragmentation as the sum of all transitions from a given sleep stage, relative to the total number of epochs in that sleep stage in the conventional sleep scoring (Figure 3). Namely, there was a higher fragmentation in W and N1 stage bouts during CSR and after ASD, respectively, and a lower fragmentation of stage N3 bouts after ASD and during CSR, which was again more pronounced after ASD. Fragmentation of stage R bouts was not significantly changed in either condition.

**Figure 3:**
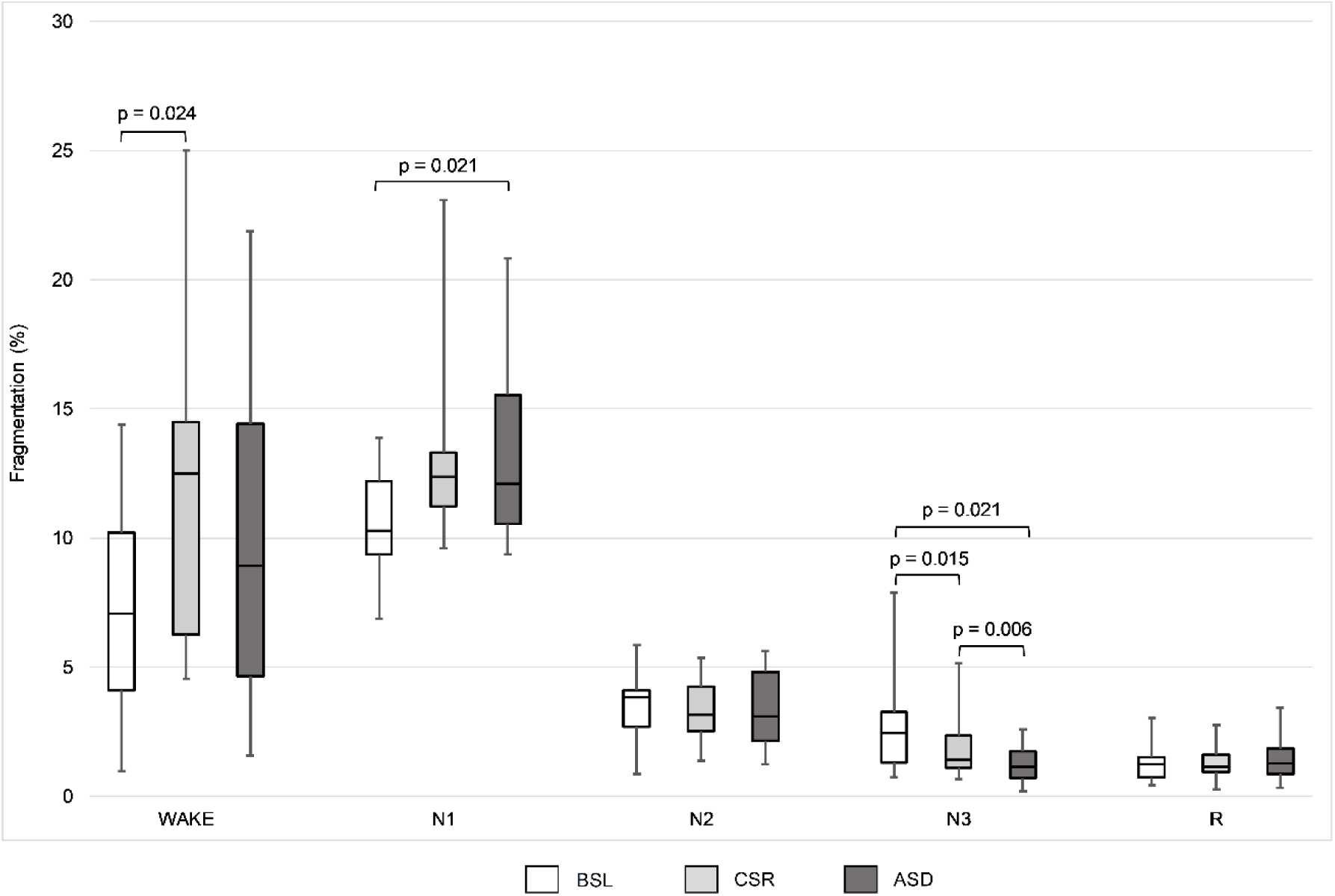
Behavioral state fragmentation. Box plots indicate medians (horizontal line), upper and lower quartiles (box), and extrema (whiskers) with corresponding p- resulting from the Wilcoxon signed-rank tests. Bonferroni correction was applied to all p-values. BSL: Baseline sleep; CSR: Chronic sleep restriction; ASD: Acute sleep deprivation; N1, N2, N3, and R refer to sleep stages as defined in the AASM Scoring Manual [36].

We further evaluated the number of transitional periods between the different sleep stages with a two-dimensional probability density approach in the SSA (Figure 4). There was no change in the total number of transitions in either condition, but significantly more of these referred to transitions between stages N2/N3 and stage R sleep after ASD. Furthermore, compared to CSR there were less transitional periods between stage R and stage W after ASD, while CSR did not result in any significant alterations of transitional periods compared to BSL.

**Figure 4:**
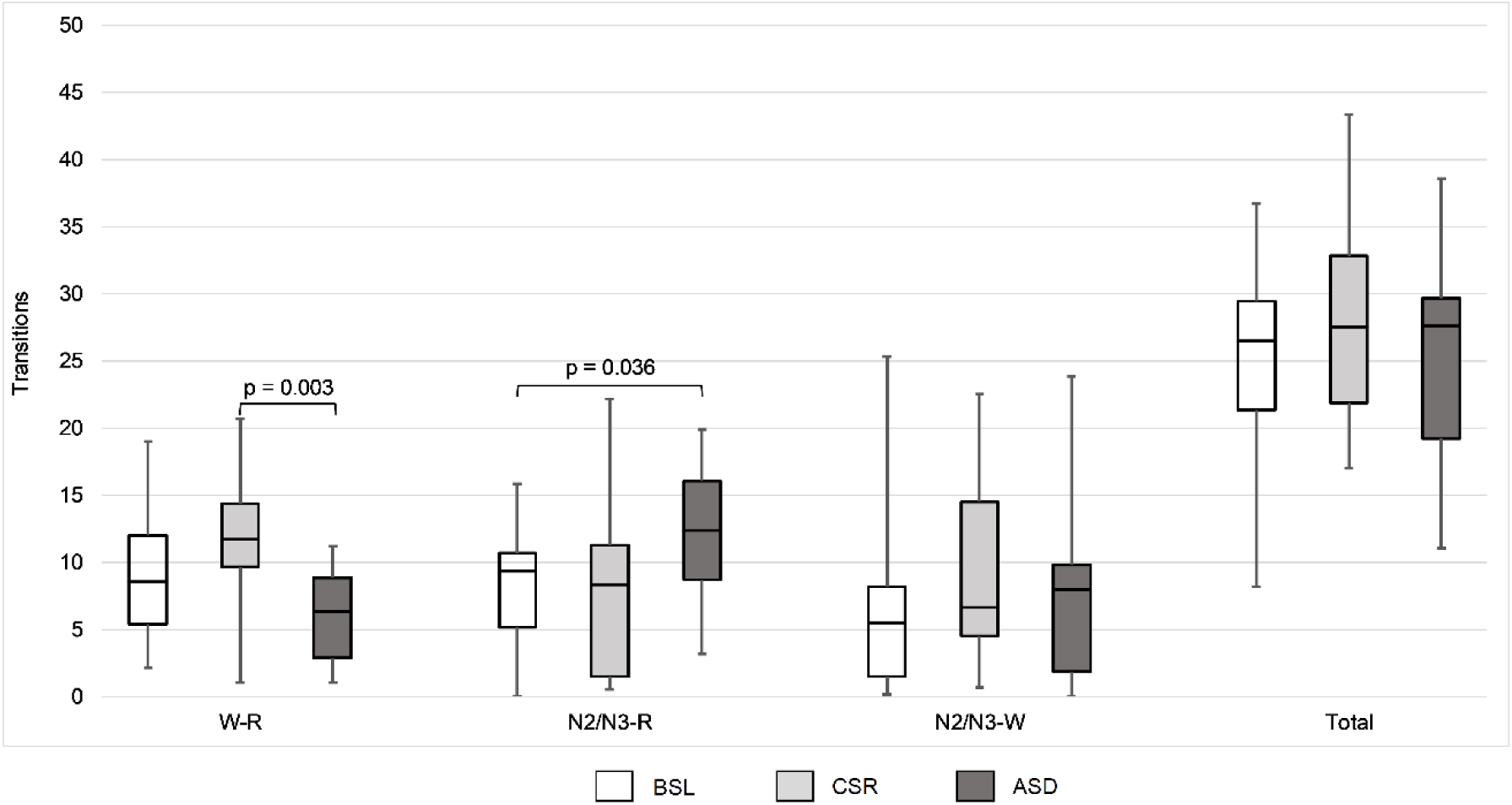
Analysis of transitional periods between the different behavioral states. Box plots indicate medians (horizontal line), upper and lower quartiles (box), and extrema (whiskers) with corresponding p-resulting from the Wilcoxon signed-rank tests. Bonferroni correction was applied to all p-values. BSL: Baseline sleep; CSR: Chronic sleep restriction; ASD: Acute sleep deprivation; N1, N2, N3, and R refer to sleep stages as defined in the AASM Scoring Manual [36].

## Discussion

In this explorative analysis, we found that experimentally induced sleep loss leads to changes in sleep characteristics that are quantifiable by SSA. The observed changes were in line with the distinct increases in SWS and REMS propensity that are expected to result from total and partial sleep loss. Of note, the analysis of conventional sleep architecture measures confirmed this assumption in our data set, with a significant increase of stage R sleep only observed during CSR and a significant increase of stage N3 sleep only evident after ASD. Thus, our findings indicate that the distinct effects of ASD and CSR are not only evident in conventional sleep architecture and spectral EEG characteristics, but are also reflected in quantitative measures of behavioral state boundaries and sleep dynamics as derived from SSA. To the best of our knowledge, this is the first within-subject comparison of behavioral state boundaries and sleep dynamics as assessed by SSA under different conditions.

Compared with conventional sleep stage scoring, SSA provides more differentiated information on spectral relations between behavioral states and their temporal stability. The analysis of stable states captures variability within sleep stages, whereas cluster-based measures describe how strongly behavioral states differ from each other in spectral terms. This may help to explain changes in conventional sleep measures such as sleep architecture, sleep latency, and transitions. Importantly, the current findings should not be interpreted as revealing novel mechanisms of sleep regulation. Rather, they indicate that SSA-derived measures are sensitive to state dependent changes in sleep organization under different homeostatic conditions. Therefore, SSA may complement conventional sleep analyses by quantifying not only the amount of a sleep stage, but also its relation to neighboring states and its stability over time.

As anticipated, we observed alterations in the spectral power composition of N2 and N3 sleep after ASD, indicated by displacements of stage N2 and N3 centroids. These affected primarily the y-dimension, mainly based on a shift in the frequency band of 8.6 Hz to 19.3 Hz and led to a significantly decreased distance between stage R and stage N2 centroids. This shift aligns with a reduction in sigma activity, which has been previously described as NREM sleep-specific changes following ASD in earlier studies.[5] Noteworthy, when considering the x- and y-dimension separately, point densities of N2/N3 and stage R revealed a higher peak-to-peak distance and a lower overlap area in x-direction, but a lower peak-to-peak distance and an increased overlap in y-direction. Thus, while N2 and stage R sleep showed an overall reduction in spectral distinction after ASD, different frequency bands were differentially affected.

Considering aspects related to sleep dynamics, we found that ASD resulted in an increased stability of stage N3 sleep, expressed by a higher ratio of stable N3 epochs compared to BSL and CSR. This is well in line with the well-known increase in sleep depth and intensity after ASD.[4, 5] Interestingly, there was no change in the total number of transitional epochs after ASD. However, we found that the number of transitions was significantly higher between stages N2/N3 and stage R than during BSL. This may be explained by the reduction in state boundaries as observed in the cluster analysis and may reflect an additional dynamic feature captured by SSA that complements the conventional findings.

Some, but not all of the findings we observed with regard to stages N3 and/or N2 after ASD were also found during CSR. Nevertheless, even if present also during CSR, these effects were mostly less pronounced than after ASD. This is well in line with previous reports that CSR also challenges SWS homeostasis, but to a much lesser extent than ASD, as large parts of SWS are preserved when curtailing sleep.[10, 13, 15] However, our explorative analysis demonstrated a clear displacement of the stage R cluster, which is in line with previously reported drifts in frequency power composition of REMS following CSR, including a decrease in alpha activity[15, 21]. The altered frequency composition of REMS resulted in a reduced distance between stage R and stage W cluster centroids after CSR, accompanied by a shortened peak-to-peak distance and an enlarged overlap area of the point density functions of stage W and stage R in x-direction. The point density analysis therefore suggests not only a shift of the stage R cluster, but also a lower separation of the distributions of stage R and stage W in the corresponding dimension. This increased assimilation reflects increased spectral similarities of REMS and wakefulness during CSR. However, this interpretation is limited by the comparatively small number of wake epochs during CSR.

Intriguingly, CSR did not alter the stability of stage R sleep. Furthermore, despite the decreased spectral state boundaries between stage R and stage W, there was no significant change in the number of transitional episodes between these states compared to BSL. These observations fit well with the previously made assumption, that alterations in state boundaries and sleep dynamics as state stability are two distinct aspects of sleep structure that may be affected independently [18, 19]. Accordingly, SSA may provide additional information beyond conventional sleep stage proportions by showing whether a behavioral state mainly changes in amount, stability, or spectral distinction from neighboring states.

The current findings suggest that state space characteristics may inform about prior sleep history, which may be of importance when assessing sleep in a clinical context. In our analysis, behavioral state boundaries and sleep dynamics changed within the same healthy subjects depending on whether prior sleep had been normal, chronically restricted, or acutely deprived. This may be especially relevant when interpreting polysomnographic recordings in populations with excessive daytime sleepiness.

SSA alterations with regard to REMS have previously been described as distinctive characteristics of sleep in patients suffering from narcolepsy. Past work showed an increased imbalance between stage R and stage W, indicated by erratic transitions between wakefulness and REMS[22] and through a higher proportion of power in the alpha frequency band in stage R sleep.[23] A recent SSA suggests that a higher similarity in spectral characteristics of the two behavioral states together with a globally impaired behavioral state stability in narcolepsy patients compared to healthy controls, may lead to the well-known pathological instability between sleep stages and wakefulness.[18]

Interestingly, we also observed a reduction in spectral distinction between stages R and W during CSR, but no evident reduction in state stability. Indeed, the lacking combination of these two aspects may be the key difference to pathological alterations as observed in narcolepsy. However, since we found some qualitatively comparable changes already after such a short-term sleep manipulation, the question arises to what degree the alterations found in narcolepsy would also be evident when comparing to a clinical relevant form of chronic sleep loss, namely in the insufficient sleep syndrome. The insufficient sleep syndrome resembles CSR and represents the most common cause for excessive daytime sleepiness,[24] evolves through self-induced sleep restriction,[25] and affects a significant portion of the population.[26–28] Actually, one could hypothesize that some common alterations in state boundaries and sleep dynamics could underlie the similarities in morphology and symptomatology between narcolepsy and insufficient sleep syndrome. [29, 30] That is for example, the occurrence of sleep onset REMS in the multiple sleep latency test [29–31] and low REMS latencies in nocturnal polysomnography. However, the distinction between these conditions with regard to sleep state boundaries and sleep dynamics remains to be investigated.

While the current analysis was restricted to young, healthy, male subjects, some cautious considerations regarding other populations may be derived from the existing literature. Previous studies suggest that female subjects may show differences in sleep EEG activity compared to male subjects, including differences in spectral power during NREM and REM sleep [32, 33]. Thus, when applying the SSA to female subjects, differences in absolute cluster positions and possibly also in cluster separation may be expected. With aging, the well described reduction in SWS and slow wave activity, together with increased sleep fragmentation, may be expected to translate into less consolidated N2/N3 sleep, a lower proportion of stable N3 epochs, and more trajectories involving stage W [34, 35]. In clinical populations, the expected alterations would likely depend on the predominant type of sleep dysregulation. In narcolepsy, previous SSA analysis already demonstrated overlap of stage R and stage W together with increased transitions between these states [18], whereas in Parkinson’s disease reduced sleep-wake dynamics have been described [19]. Thus, future studies are needed to determine to what extent the present approach may also help to characterize sex-, age-, and disease-related alterations in sleep structure and sleep dynamics.

There are limitations of the current analysis, which need to be taken into consideration. For facilitating comparison between the CSR, ASD and BSL nights, we had to shorten the BSL night and the recovery night after ASD to the 5 hours duration of the CSR night. Therefore, our analysis cannot make any statements on the last part of a normal sleep period, that may entail more REMS, but also more intrusions of brief episodes of wakefulness [16]. This could of course alter overall cluster compositions and/or sleep dynamic measures. In addition, the shortened analysis period together with the high sleep pressure in the sleep deprivation conditions resulted in a comparatively low number of wake epochs, and to a lesser extent N1 epochs, particularly in the stable state analysis. Therefore, measures involving wakefulness need to be interpreted with caution. At the same time, the low amount of wakefulness is likely related to the experimental conditions and may reflect that wake episodes within the analyzed period mainly occurred as brief arousal-related intrusions under increased sleep pressure. While these aspects pose limitations from a scientific point of view, the situation may be comparable to some clinical contexts. Nevertheless, the presented results are restricted to a healthy, young, male population and further investigations are needed for generalization, particularly in female subjects, older individuals, and clinical populations with altered sleep regulation or sleep structure.

## Conclusion

Taken together, the results of our explorative study suggest that the differential increases in REMS and SWS propensity are reflected in distinct changes of behavioral state boundaries and sleep dynamics as derived from SSA. Furthermore, our findings indicate that these sleep structure characteristics are state dependent and that such measures may complement conventional approaches by providing quantitative information on the spectral differentiation and stability of behavioral states. Future studies are needed to determine whether these measures may also be useful for characterizing sleep structure alterations in larger samples and in clinical populations.

## Acknowledgement

We thank Rositsa Poryazova and Esther Werth for supervising the study conduction, Caroline Lustenberger and Reto Huber for support in sleep data analysis setup and Janina Leemann, Felicitas Gilgen, Cornelia Wettstein, Eszter Montvai, Vanessa Sennrich, Manuel Bürgi, Matthias Storz, Laura Kopacsi, Jenni Saarto, Jan Steiner, Alla Mühlebach, Annina Bieri, Mara M. Suter and Susanne Kanzler for help in data acquisition.

The study was funded by the Clinical Research Priority Program (CRPP) Sleep and Health of the University of Zurich, the Olga Mayenfisch Foundation, and the Gender Equality Action Plan of the University of Zurich Filling the Gap.

## Data availability statement

Data available on reasonable request.

## Author contributions

Conceptualization: AM, CB, LI, SR; Methodology: AM, LI, SR; Software: SR, LI; Validation: LI, AM, CB; Formal analysis: SR; Investigation: SR, ZB; Resources: AM, CB; Data curation: ZB, SR; Writing – original draft: SR, AM, LI; Writing – review & editing: AM, LI, CB, SR; Visualization: SR; Supervision: AM, LI, CB

